# Single cell RNA-sequencing reveals GINIP-expressing neurons as the main targets of focused ultrasound

**DOI:** 10.1101/2024.07.17.604026

**Authors:** Elena Brunet, Thibaud Parpaite, Sungjae Yoo, Eric Debieu, Khaled Metwally, Serge Mensah, Pascale Malapert, Andrew Saurin, Olivier Macherey, Emilie Franceschini, Aziz Moqrich

## Abstract

Dorsal root ganglion (DRG) neurons have a wide range of functions, including touch, pain and itch. These neurons have emerged as promising targets for non-invasive focused ultrasound (FUS) neuromodulation. However, our knowledge of the molecular and physical mechanisms underlying FUS-evoked responses in DRG neurons is limited. Here, we investigate the neuromodulatory capabilities of FUS in cultured DRG neurons in combination with calcium imaging. We find that a 20-MHz FUS burst of 1-ms duration at an acoustic pressure of 5 MPa elicited calcium responses in 52% of DRG neurons. Single-cell RNA sequencing reveals that the majority of FUS-sensitive neurons belong to three subsets of DRG neurons; C-LTMRs, the MRGPRD-expressing C-HTMRs and A6-LTMRs. FUS excites all these neuronal subtypes by membrane deformation, suggesting a mechanism mediated by mechanosensitive ion channels. Our results identify FUS parameters that activate distinct subsets of DRG neurons and open new avenues for using FUS stimulation to modulate DRG neuron function.

## INTRODUCTION

Sensory neurons of the dorsal root ganglia (DRG) play a crucial role in transmitting sensory information from the periphery (such as skin, muscles and organs) to the central nervous system (CNS) for further processing and interpretation. These neurons exhibit a wide range of molecular, morphological and physiological properties that underlie their selective responses to various environmental stimuli, including temperature, pressure, itch and pain. Pathological conditions such as inflammation and nerve injury can sensitise DRG neurons, leading to an exaggerated response to noxious stimuli (hyperalgesia) and abnormal activation to innocuous stimuli (allodynia), often resulting in chronic pain (1). Chronic pain is a devastating condition that affects more than 30% of people worldwide (2). Opioids have long been used as potent analgesics and are considered the mainstay of chronic pain management. However, the opioid epidemic, characterised by an increase in opioid-related deaths and overdoses, has been recognised as a significant public health crisis (3). Therefore, efforts to develop alternative strategies for the treatment of chronic pain while avoiding harmful side effects should be encouraged.

Focused ultrasound (FUS) is a non-invasive medical technology that uses ultrasound to target specific tissues for therapeutic or diagnostic purposes. Transcranial FUS has been used to treat pain by targeting specific brain circuits. FUS stimulation of the periaqueductal grey (PAG) effectively suppresses formalin-evoked pain in rats (4), and stimulation of the primary somatosensory cortex (S1) significantly attenuates heat pain sensitivity in wild type mice, and modulates injury-induced thermal and mechanical hyperalgesia in a mouse model of sickle cell disease (5). Although very encouraging, these attempts mainly focused on modulation of the activity of central but not peripheral nervous system. In a recent elegant study, Hoffman and colleagues combined FUS stimulation and electrophysiology in a mouse skin-nerve preparation and found that stimulating neuronal receptive fields with high-intensity, millisecond FUS pulses triggered action potential in nearly all recorded fibers partly in a PIEZO2-dependent manner (6). This pioneering study provides the first demonstration that primary sensory neurons can be effectively and reliably modulated by FUS. Importantly, the study also showed that low-threshold mechanoreceptors exhibited a much-pronounced FUS sensitivity than C-nociceptors, revealing a potential functional heterogeneity in sensory neurons in response to FUS stimulation. In this study, we combined FUS stimulation with calcium imaging and single cell RNA sequencing to confirm that DRG neurons indeed respond to FUS stimulation and to provide a comprehensive transcriptomic analysis of FUS sensitive and insensitive neurons. We found that a 20-MHz FUS stimulus for 1-ms duration at an acoustic pressure of 5 MPa triggered calcium responses in 52% of DRG neurons. Mapping the transcriptional profiles onto previously identified sensory neuron subtypes shows that FUS-sensitive neurons can be found in nearly all categories of neurons, with a higher prevalence in GINIP-expressing neurons. Ultra-fast camera monitoring of FUS-mediated membrane displacement showed that cell deformation increases linearly with increasing acoustic pressure or duration, and that this phenomenon was more pronounced in GINIP^-^ neurons.

## RESULTS

### Focused ultrasound activates DRG neurons

Simultaneous calcium imaging and FUS stimulation were used to monitor neuronal responses in cultured primary sensory neurons isolated from DRGs of R26^GCaMP6S/+^::Advillin^Cre/+^ mice (Fig. 1*A*). Ultrasound was delivered using a transducer with a central frequency of 20 MHz and with focal distance of 12.7 mm. The transducer was mounted on a 3D motorized micromanipulator with a tilt angle of about 20° to reduce standing wave formation. Prior to FUS stimulation, the US beam was positioned at the center of the optical microscope’s field of view. To achieve this, the transducer was first operated in pulse-echo mode and moved laterally to maximize the ultrasonic echo signal from an isolated microsphere placed at the center of the field of view. For the FUS stimulation, a fundamental sinusoidal frequency of 20 MHz was used. The sinusoidal signal was produced by a function generator and amplified with a power amplifier. The resulting US beam in free-field had a focal diameter of 224-µm, as measured by the -6 dB beam width (Fig. 1*A*).

**Figure 1.**
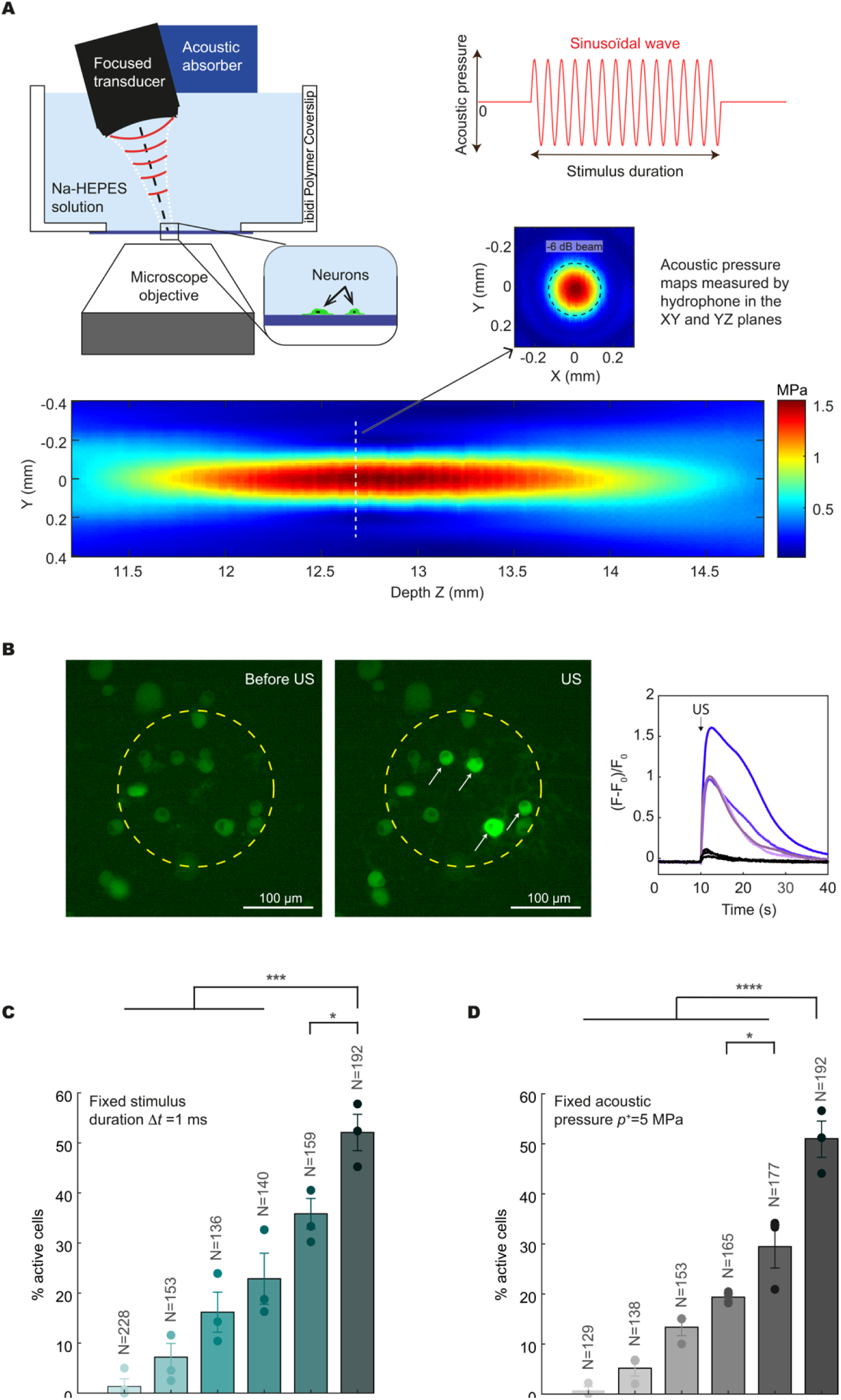
FUS at 20 MHz activates cultured DRG neurons. (*A*) Schematics of the FUS stimulation setup and the ultrasound waveform applied to neurons. Ultrasounds are delivered to GCaMP6s DRG neurons cultured on a polymer dish, while the neural responses are recorded by calcium imaging. Hydrophone measurements of the ultrasound beam profiles emitted by the 20 MHz transducer in the X-Y and X-Z planes. (*B*) Examples of images of GCaMP6s fluorescence before and during FUS stimulation for the stimulus [*p^+^*=5 MPa, Δ*t*=1 ms], and corresponding calcium responses of DRG neurons within the -6 dB beam area. (*C*) Percentage of FUS-sensitive neurons as a function of acoustic pressure (n = 3 independent experiments, one-way ANOVA, *** p < 0.0001, * p = 0.0445) and (*D*) stimulus duration (n = 3 independent experiments, one-way ANOVA, *** p < 0.0001, * p = 0.0391). Bar graph values represent mean ± SEM. N is the total number of analyzed cells.

Dissociated DRG neurons were plated on an imaging dish with a polymer coverslip coated with laminin, allowed to adhere overnight, and then sonicated with FUS. To identify the US pulse parameters for which DRG neurons are activated, cells were stimulated by varying acoustic positive pressure (*p^+^*=4-5 MPa, corresponding to Spatial Peak Pulse Average intensity I_sppa_=346-536 W/cm^2^) and stimulus durations (Ll*t*=0.1-1 ms) (Table S1). DRG neurons could be detached by FUS at high acoustic pressures (>5 MPa for 1 ms stimulus) or longer stimulus durations (>1 ms for 5 MPa stimulus), limiting our study to these maximum values. In the following, the stimulus consisting of a positive pressure of 5 MPa and a duration of 1 ms is referred to as the “optimal FUS stimulus”. Each cell was subjected to a single FUS stimulus to avoid potential cumulative effects. The percentage of FUS-sensitive neurons were computed by considering only the DRG neurons within the -6 dB beam area (Fig. 1*B*). Within the parameter ranges tested, the percentage of FUS-sensitive neurons increased both with increasing acoustic pressure and with stimulus duration (Fig. 1*C* and *D*). The optimal FUS stimulus activated 52% of DRG neurons (N=192 cells). To ensure the integrity of the cultured neurons, the last US stimulation session for each coverslip was followed by a bath application of potassium chloride (KCl) (*SI Appendix*, Fig. S1*A*). We observed three different calcium kinetics on the FUS-sensitive neurons: rapid (T1), intermediate (T2) and slow (T3) (*SI Appendix*, Fig. S1 *B* and *C*). Rapid calcium transients seem to dominate at low acoustic pressures, while the percentage of intermediate and slow calcium kinetics increases with acoustic pressures (*SI Appendix*, Fig. S1*B*). Acoustic pressure seems to influence the calcium kinetics more than stimulus duration, since no clear relationship can be observed between stimulus duration and the types of calcium kinetics (*SI Appendix*, Fig. S1*C*).

### FUS triggers calcium responses preferentially in GINIP-expressing neurons

Given that FUS stimulation activates only a subset of DRG neurons, we sought to determine the molecular identities of FUS-sensitive and FUS-insensitive neurons. We combined calcium imaging with FUS stimulation and single cell RNA sequencing (sc-RNA-seq) (Fig. 2*A*). Cultured DRGs neurons from R26^GCaMP6S/+^::Advillin^Cre/+^ mice were sonicated with two consecutive optimal FUS stimulations with a two minute interval between each stimulation. FUS-sensitive neurons were defined as the neurons responding to the first stimulus, while FUS-insensitive neurons were those that did not respond to any of the two stimuli. Responsive and non-responsive neurons were individually picked using a patch-clamp pipette and the corresponding cDNA libraries were generated for RNA-seq. Using this approach, out of the 120 picked neurons, 57 passed all the quality controls required for sequencing, including 45 FUS-sensitive and 12 FUS-insensitive neurons. To identify to which categories of neurons the FUS-sensitive and FUS-insensitive neurons belonged, we performed the same analysis as Parpaite and colleagues (7). We merged our data with the publically available datasets generated by Zeisel and colleagues using fast mutual nearest neighbors (fastMNN) correction (8,9). We found that FUS-sensitive neurons spread across all previously identified neuronal subsets except for the TRPM8 population (Fig. 2 *B* and *C*, and *SI appendix*, Fig. S2). However, out of the 45 FUS sensitive neurons, 56% (25 out of the 45) belong to the TH-expressing C-LTMRs (12) and the MRGPRD-expressing NP1 (13), both of which express the GINIP encoding gene *Phf24* (9) (Fig. 2*D*). In line with this, FUS-insensitive neurons were excluded from five out of the nine neuronal subsets: the TH^+^ C-LTMRS, the MRGPRD-expressing NP1, the MRGPRA3-expressing NP2, the Sst-expressing NP3 and the TrkB-expressing NF1 neurons (Fig. 2*D*).

**Figure 2.**
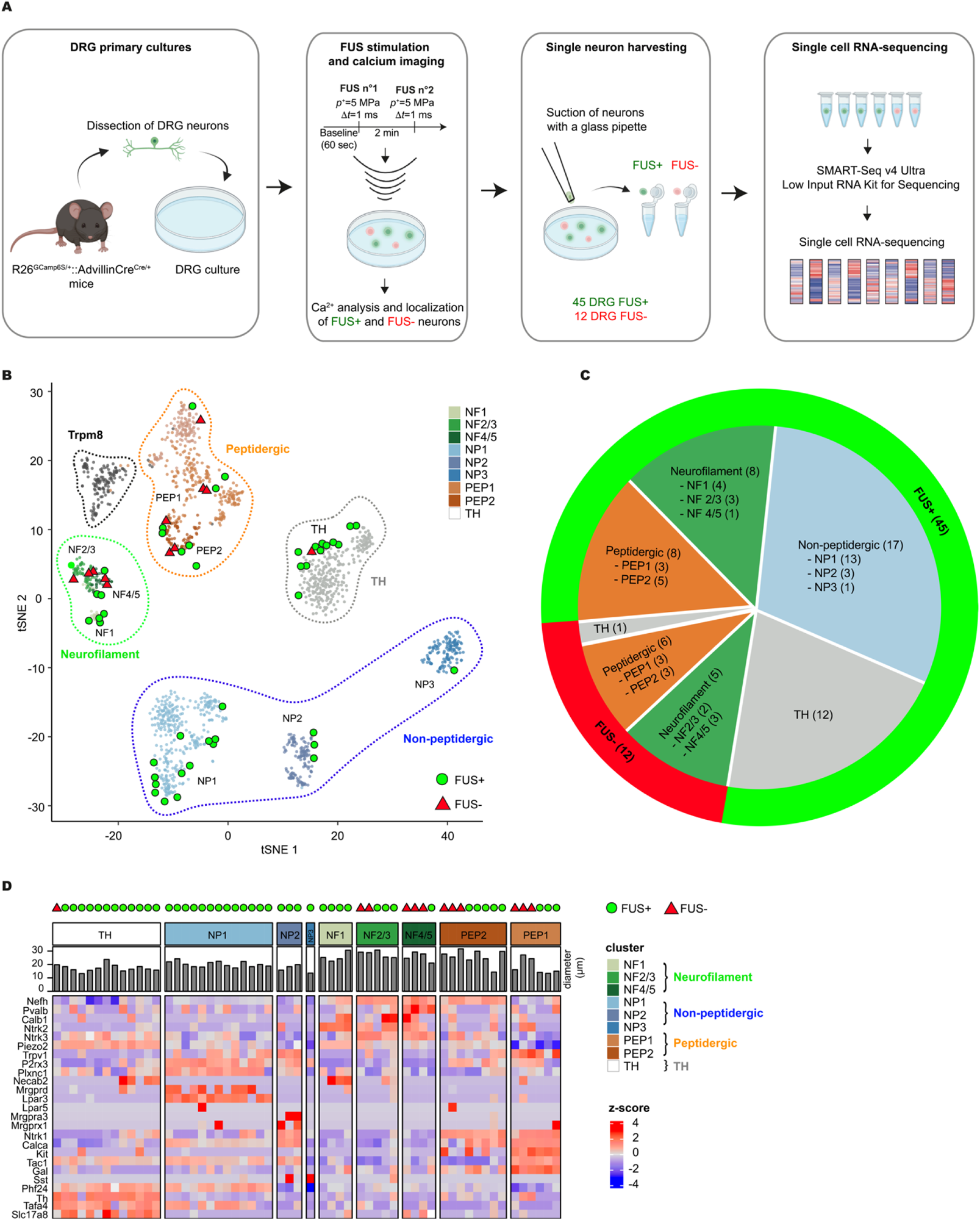
Expression profiling of FUS-sensitive neurons. (*A*) Schematic representation of the methodology used in this study. (*B*) Visualization using t-SNE embedding of all cells colored by cluster identity. Red triangles represent FUS-insensitive neurons and green circles represent FUS-sensitive neurons from sc-RNAseq. The small light circles represent the scRNA-seq neurons from the study of Zeisel and colleagues. Main populations of DRG neurons are surrounded by dashed lines. (*C*) Quantification of the number of FUS-sensitive and FUS-insensitive neurons from sc-RNAseq. (*D*) Heatmap of marker gene expression of sequenced FUS-sensitive and FUS insensitive neurons grouped by population (NF, neurofilament; NP, non-peptidergic; TH, tyrosine hydroxylase; PEP, peptidergic).

### Molecular determinants underlying DRG neurons FUS sensitivity

A detailed analysis of the calcium transients in the sequenced FUS-sensitive neurons revealed three distinct response profiles with regard to their inactivation kinetics: rapid (8 out of 45 neurons), intermediate (15 out 45 neurons) and slow (21 out 45 neurons) (Fig. 3*A*). Neurons belonging to the NF population respond to FUS stimulation with intermediate or slow inactivation kinetics, whereas those belonging to the NP, PEP and TH subsets exhibit the three types of kinetics (Fig. 3*B*). To assess the molecular determinants underlying FUS sensitivity and the differences in calcium decay time, we performed several comparisons using our single-cell transcriptomic data. First, we analyzed the transcriptomic differences between FUS-sensitive and -insensitive neurons. We performed WGCNA to analyze 885 differentially expressed genes between FUS-sensitive and -insensitive neurons (see *Supplementary Dataset Table E2*). We detected three significant gene modules, with two up-regulated modules (orange and brown) and one down-regulated module (pink) (Fig. 3*C*). The 284 genes in the orange module are highly expressed in FUS-sensitive neurons with a more pronounced upregulation in the subclass characterized by a rapid inactivation kinetic (Fig. 3*D* and *Supplementary Dataset Table E3*). Gene ontology (GO) analysis of this module showed the enrichment for protein localization in cell associated terms such as localization to plasma membrane or establishment of protein localization to the mitochondrion. The 179 genes in the brown module are also upregulated in FUS-sensitive neurons but have the same expression level regardless of their calcium response kinetics. Among these genes, we have an enrichment in potassium channel associated genes such as Kcng2, Kcnip4, Kcnk13, Kcnmb1 and Kcnmb4 as well as genes involved in cellular calcium ion homeostasis and pain response. By contrast, the 238 genes in the pink module were preferentially expressed in FUS-insensitive neurons, and showed an enrichment in genes associated with collagen fibril organization and cell-matrix adhesion (Fig. 3*D*).

**Figure 3.**
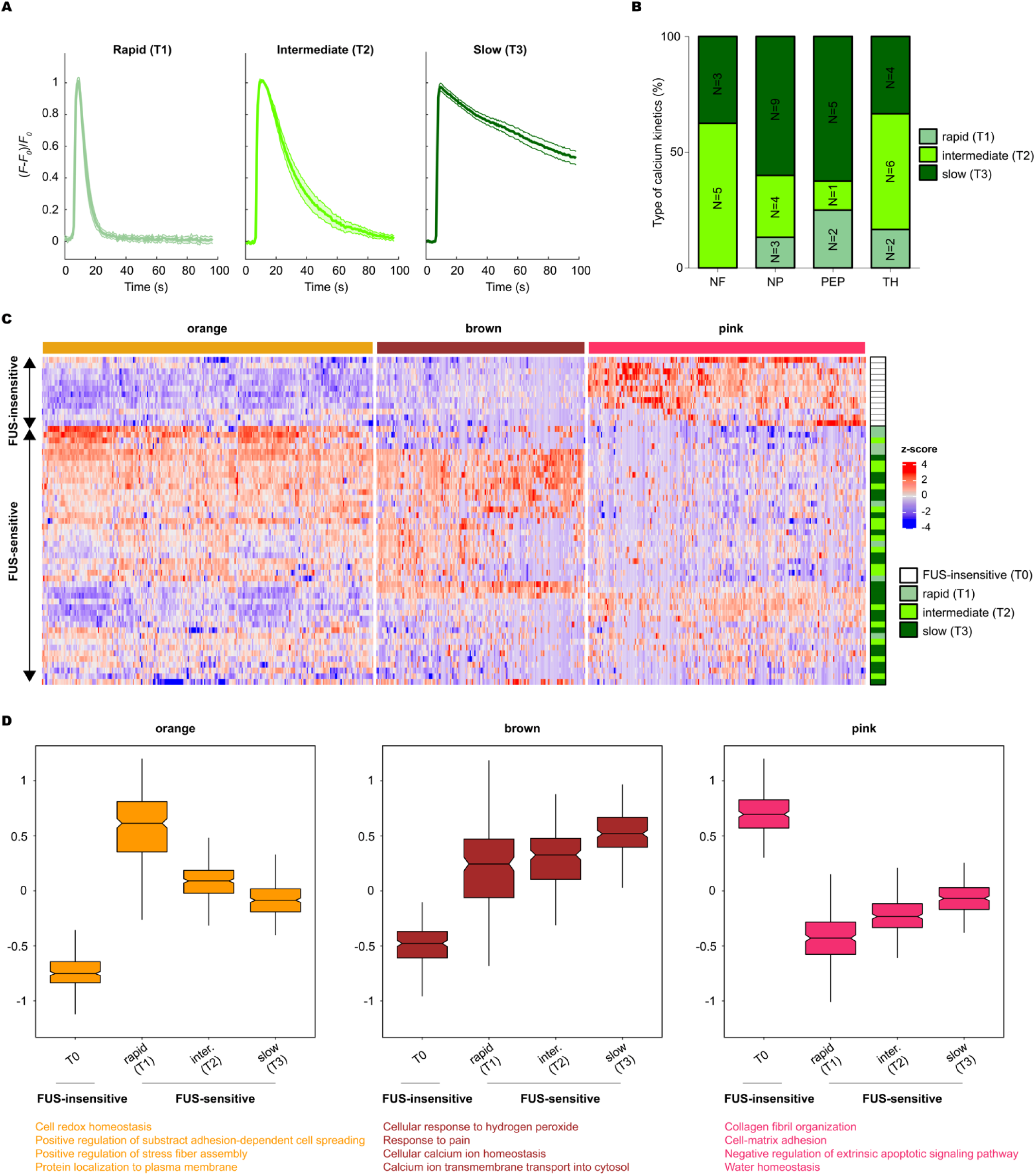
Associated genes with calcium kinetics of FUS-sensitive neurons. (*A*) Normalized mean calcium responses of the 3 types of kinetics observed in sequenced FUS-sensitive neurons (slow, intermediate and fast inactivation). The mean trace is shown in light, medium and dark green and the SEM is shaded. (*B*) Proportion of the 3 types of calcium kinetics of scRNA-seq FUS-sensitive neurons in each DRG population (NF, neurofilament; NP, non-peptidergic; PEP, peptidergic; TH, tyrosine hydroxylase). N is the number of cells. (*C*) Heatmap showing relative expression of genes in three gene modules identified by WGCNA in FUS-sensitive and FUS-insensitive neurons. (*D*) Boxplots showing expression patterns (scaled log2 TMM) of differentially expressed genes in each WGCNA module. Representative GO (gene ontology) terms of each significantly regulated module are listed below.

### FUS excites GINIP-expressing neurons via membrane deformation

The sc-RNAseq analysis shows that 56% of the FUS-sensitive neurons belong to the GINIP^+^ neuron class, which consists of the TH-expressing C-LTMRs and the MRGPRD-expressing NP1 (Fig. 4*A*). To confirm this result, calcium imaging was performed on DRG neurons from GINIP^flx/+::^Advillin^Cre^ mice (hereafter called GINIP^mCherry^ mice), which allow expression of mCherry from the *ginip* locus. For both GINIP^+^ and GINIP^-^ neuron types, the percentage of FUS-sensitive neurons was calculated using the same optimal FUS stimulus [*p*^+^=5 MPa, Δ*t*=1 ms] as used for the sequenced neurons. In this mouse model, FUS stimulation elicited calcium transients in 59% of the DRG neurons (grey bars) (Fig. 4*B*), which matched the 52% of FUS-sensitive neurons observed in the R26^GCaMP6S/+^::Advillin^Cre/+^ mice and shown in Figure 1*D*. To confirm that GINIP^+^ neurons are more sensitive to FUS stimulation than other types of neurons, we performed calcium-imaging experiments on mCherry^+^ and mCherry^-^ neurons. Consistent with our RNA-seq data, we found that an acoustic pressure of 5 MPa for 1 ms elicited a calcium transient in 72% of mCherry^+^ neurons (green bars), whereas the same stimulation excited only 47% of mCherry^-^ neurons (pink bars) (Fig. 4*B*). Among the mCherry^+^ neurons, a bath application of isolectin IB4 allowed the identification of two distinct subsets of neurons: the NP1 mCherry^+^-IB4^+^ neurons and the C-LTMRs mCherry^+^-IB4^-^ neurons. Using this approach, we found that FUS stimulation elicited calcium responses in 84% of C-LTMRs (white bars) and 67% of MRGPRD+ NP1 neurons (sky-blue bars) (Fig. 4*B*).

**Figure 4.**
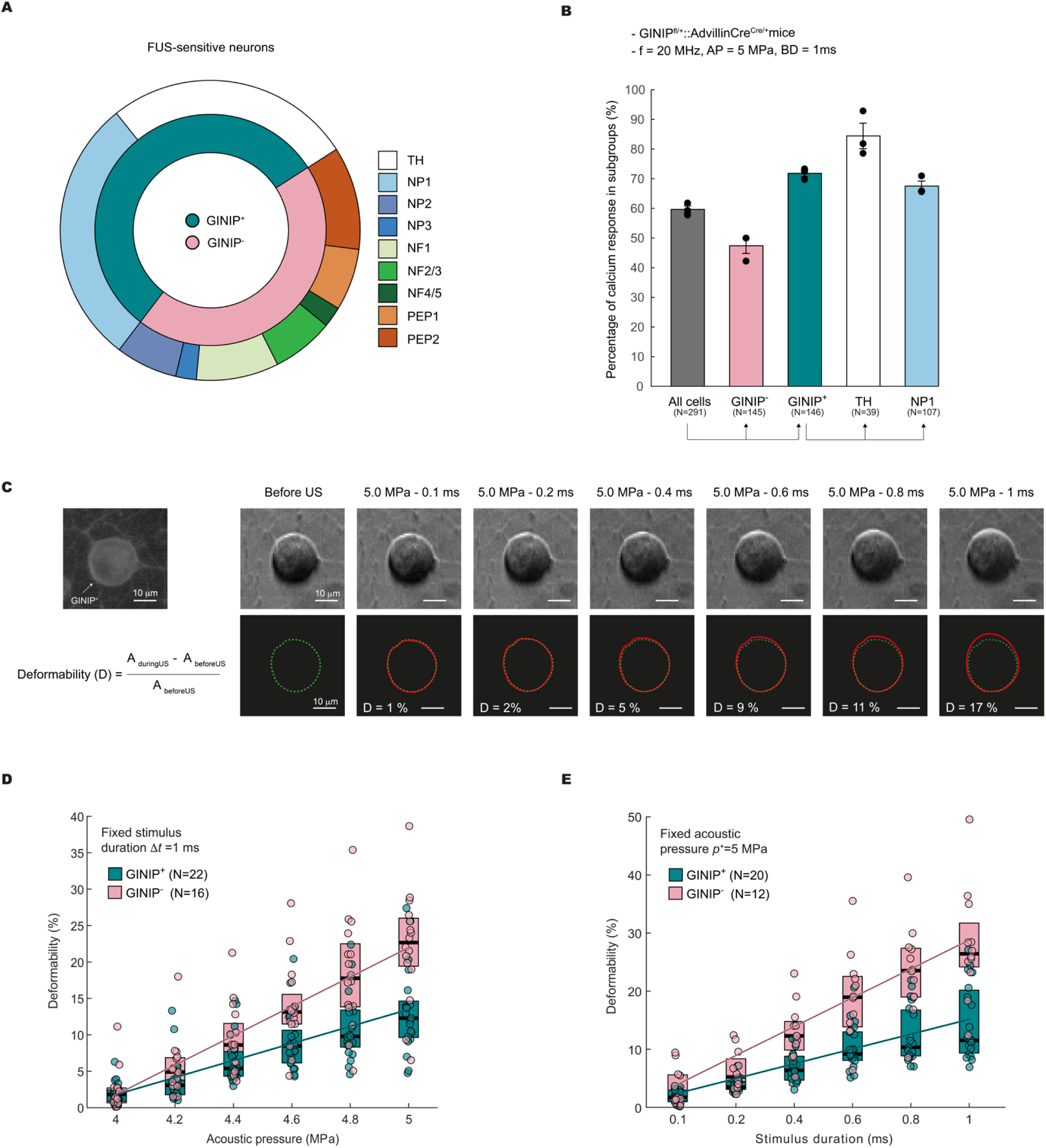
FUS excites GINIP-expressing neurons via membrane deformation. (*A*) Proportion of neurons of different classes among all FUS-sensitive neurons. GINIP-expressing neurons represent 56% of all DRG neurons responding to FUS. (*B*) Percentages of responding cells in GINIP^-^ and GINIP^+^ neurons, and in GINIP^+^ subpopulations (TH and MRGPRD). (*C*) Example of deformability of a GINIP^+^ neuron. Comparison between the cell projected area before and during FUS stimulation. The green and red lines represent the contour of the cell before and during FUS stimulation, respectively. (*D*) Boxplots showing the deformation of GINIP^+^ and GINIP^-^ neurons as a function of acoustic pressure with a fixed stimulus duration of Δ*t*=1 ms. The solid lines represent the output of a linear mixed model with random factors (intercept and slope) and fixed factors (acoustic pressure and neuron type: GINIP^+^ and GINIP^-^). N is the total number of analyzed cells. (*E*) Same as (*D*) for the stimulus duration with a fixed acoustic pressure of *p^+^*=5 MPa.

Given the high acoustic pressure involved in our experiments, we hypothesized that FUS-mediated calcium transients are likely due to membrane deformation. The mechanical effects in our set-up may be caused by acoustic radiation forces or hydrodynamic forces induced by acoustic streaming, or a combination thereof. Both acoustic radiation and acoustic streaming are second order nonlinear acoustic effects. Acoustic radiation force is the net force experienced by the cell resulting from the nonlinear interaction between the cell and the ultrasonic wave. Acoustic streaming is the steady fluid flow generated by the attenuation of the ultrasonic wave, resulting in hydrodynamic forces along the cell surfaces. We examined whether the DRG neurons were deformed due to the US-induced mechanical forces. To this end, we used GINIP^mCherry^ to monitor FUS-induced deformability in a homogenous population of neurons. The cellular deformation was monitored using an ultra-fast camera (Photron FASTCAM, 100000 fps) for the same US stimuli as used in the quantification of FUS-sensitive neurons as a function of duration and pressure (Fig. 1*C* and *D*). The cellular deformation was computed as the ratio between the change in the projected area of the cell and the projected area of the undeformed cell before US stimulation. Figure 4*C* illustrates how the projected area became larger as the stimulus duration was increased. Linear mixed models were fitted to the deformation data, as detailed in the Methods section. Cell deformation monotonically increased with increases in acoustic pressure (Fig. 4*D*) or stimulus duration (Fig. 4*E*), as demonstrated by significant main effects of these two factors compared to the random intercept-only models (χ²(1)=289, p<0.0001 for pressure χ²(1)=226, p<0.0001 for duration). There was also a significant interaction of each of these two factors with the neuron types GINIP^+^ and GINIP^-^ (χ²(2)=69, p<0.0001 for pressure and χ²(2)=80, p<0.0001 for duration). This indicates a larger deformation for GINIP^+^ than GINIP^-^ when increasing stimulus pressure or duration (Fig. 4 *D* and *E*). Finally, the models were further improved by allowing the deformation vs. pressure or duration slope to vary (χ²(2)=142, p<0.0001 for pressure and χ²(2)=164, p<0.0001 for duration). As shown in Fig. 1 *C* and *D*, the lowest stimulus duration required to activate DRG neurons was 0.1 ms at the maximum acoustic pressure of 5 MPa, and the lowest acoustic pressure to activate DRG neurons was 4 MPa at the maximum stimulus duration of 1 ms. It is interesting to note that no cell deformation could be observed below these thresholds of 0.1 ms and 4 MPa. This suggests that cell deformation greater than a few percent is required to activate DRG neurons via mechanoreceptors.

What are the mechanisms underlying the difference in deformability between GINIP^+^ and GINIP^-^ neurons? To address this question, we interrogated our sc-RNA-seq data to see whether some gene-property relationships might be potentially different within or specific to, GINIP^+^ or GINIP^-^ neurons, reflecting a graded phenotypic difference within these cell types. To test this, we included an interaction term between gene expression and GINIP^+^ versus GINIP^-^ neuron class to assess whether the gene-property relationships (i.e. maximum amplitude of the calcium response) were different within each cell class (see Methods for further details and *Supplementary Dataset Table E5*). Interestingly, we found that the mRNA level of Kcnip4, which encodes potassium channel interacting protein 4, strongly correlates with the cell-to-cell variability in the maximum amplitude of the calcium response between GINIP^+^ and GINIP^-^ neurons (*SI Appendix*, Fig. S4). Furthermore, the mRNA level of Flot1, which plays a role in vesicle trafficking and cell morphology, correlates with the cell-to-cell variability in the inactivation kinetics of the calcium response between GINIP^+^ and GINIP^-^ neurons (*SI Appendix*, Fig. S4). This relationship is not apparent when this class of neuron is not considered, probably reflecting that a different cellular context (i.e. membrane lipid composition) influences the neuron’s response to FUS.

## DISCUSSION

### Identity of FUS-sensitive neurons

Targeted non-invasive neuromodulation offers exciting opportunities for both clinical and research applications, with the potential to treat neurological and psychiatric disorders and to improve our understanding of brain function. The use of ultrasound to activate peripheral neurons began more than 40 years ago. By stimulating the human arm with short FUS pulses, it was possible to mimic natural stimuli such as touch, temperature and pain. These different sensory modalities can be evoked at the same site on the skin by adjusting the intensity of the US stimulus, suggesting that FUS can selectively activate different types of sensory nerve fibers in the peripheral nervous system, leading to different sensory experiences (11). A recent study from the Lumpkin lab provided the first ex vivo demonstration that a brief high-intensity stimulation with FUS elicits action potentials in the peripheral endings of all types of mechanosensory neurons (6). In our study, we took a complementary approach with two ideas in mind. The first was to see if the findings of Hoffman and colleagues could be extrapolated to the cell bodies of DRG neurons. If so, having access to the RNA content of all responding cells will allow the identification of the molecular determinants underlying FUS sensitivity. We found that FUS sonication of 5 MPa for 1 ms elicited robust calcium responses in 52% of cultured DRG neurons. Single-cell RNA-sequencing revealed that FUS-sensitive neurons belong to almost all neuronal categories, with a clear bias towards five distinct neuronal subsets; the C-LTMRs, the MRGPRD-expressing mechano-nociceptors, the MRGPRA3-expressing itch-sensing neurons, the Sst-expressing NP3 neurons and the D-hair Ao-LTMRs. Our data only partially overlap with the study by Hoffman and colleagues, who showed that sonication of mouse saphenous receptive fields elicited action potentials in all myelinated and unmyelinated fibers (6). How can we explain such a difference between the two studies? The first plausible explanation is the experimental approach. We used calcium imaging on cell bodies, whereas they used electrophysiological recordings on peripheral terminals. The electrophysiological approach offers better spatial and temporal resolution compared to calcium imaging, which may not accurately capture fast neuronal events. Second, peripheral nerve endings are likely to be more sensitive to FUS stimulation than the cell bodies of DRG neurons because the molecular machinery involved in FUS sensitivity, such as ion channels, mechanosensitive proteins or cytoskeletal elements, is optimally localized in the membrane of nerve endings compared to cell bodies. As the electrophysiological recordings in the Hoffman study were performed on intact adult peripheral terminals, it is likely that the mechanotransduction pathways activated by FUS stimulation are already in place. In line with this, Yoo and colleagues showed that 12-14 days old primary murine cortical neurons are activated by FUS through a mechanical mechanism involving a gradual increase in calcium that is amplified by calcium and voltage-gated channels, resulting in burst firing activity (12).

### Physical mechanisms of FUS neurostimulation

Several mechanisms based on thermal or mechanical effects have been proposed to explain FUS neurostimulation. Numerous studies have shown that FUS can modulate neural activity without causing significant tissue heating (13). Only minimal thermal increases of less than 1°C have been reported during in vitro or in vivo experiments of FUS applications for neurostimulation (6, 12, 14–16). In our experiments, FUS can increase the temperature of the laminin-coated polymer coverslip to which DRG neurons are attached. We predict that the US stimulus with a maximum acoustic pressure of *p*^+^=5 MPa for a pulse duration of Δ*t* =1 ms should produce a temperature increase of less than 2°C (see Methods). As all our experiments were performed at room temperature (between 20 and 22°C), a temperature rise of 2°C is largely insufficient to elicit calcium responses, suggesting that the FUS responses were driven by mechanical forces.

Among the potential mechanical effects of US, bubble formation and cavitation may be involved as a FUS stimulation mechanism (17). However, the likelihood of cavitation inception decreases when using higher frequencies and lower acoustic pressure. For our maximal US stimulus parameters with peak negative pressure *p^-^*=-3.2 MPa, the mechanical index (*MI*=0.7) is below the cavitation threshold in soft tissues (*MI*>1.9). The presence of cavitation is therefore unlikely under our experimental conditions. This is consistent with the good agreement obtained between hydrophone measurements and nonlinear simulations of the 3D wavefield at the focus (*SI Appendix*, Fig. S5 *A* to *F*), showing that the nonlinearity observed in our experimental conditions are caused by the nonlinear propagation of the focused US and not by oscillating bubbles (*see Methods*).

The most likely hypothesis is that FUS-mediated neurostimulation occurs through mechanical effects on the DRG neurons, via acoustic radiation force and/or streaming-induced force, which leads to the activation of ion channels and generation of neuron action potentials. Indeed, previous *in vitro* experiments on transfected cells demonstrate that Piezo1 channel can be activated by FUS through cell membrane stress caused by acoustic fluid streaming (18, 19). Recent studies on *ex vivo* retina (16) and *in vivo* nerves (20) showed correlations between acoustic radiation force (via tissue displacement) and neural activity. Using high speed imaging during FUS stimulation, our results show that cell deformation is required to activate DRG neurons. This reinforces the hypothesis that FUS stimulation on DRG neurons is driven by cell deformation as a result of the US-induced mechanical forces. Finally, our experiment demonstrates the reduced deformation of GINIP^+^ compared to GINIP^-^ under FUS stimulation (Fig. 4 *D* and *E*). A possible explanation could be the higher Young’s modulus of the GINIP^+^ subtype, which will be further investigated in future research.

### Molecular mechanisms of FUS neurostimulation

Our study revealed that FUS-sensitive neurons responded with three different profiles based on their inactivation kinetics: rapid, intermediate, slow, with the NF population predominantly exhibiting intermediate and slow kinetics. Given that we have used a genetically encoded calcium indicator, the GCaMP6s, our data suggest that FUS stimulation likely activates different signaling pathways that lead to different amount of intracellular calcium concentrations responsible for the different calcium decay times. To underpin the mechanisms underlying FUS sensitivity and the variations in calcium decay time, single-cell RNA profiling of FUS-sensitive and FUS-insensitive neurons identified 885 differentially expressed genes that could be categorized in three significant gene modules. Two gene modules enriched in FUS-sensitive neurons and one gene module enriched in FUS-insensitive neurons. The first module, comprising 284 genes, showed high expression in FUS-sensitive neurons, particularly in the subclass characterized by rapid inactivation kinetics. GO analysis highlighted enrichment in terms related to protein localization within cells, such as plasma membrane localization and establishment of protein localization to the mitochondrion. The second one, consisting of 179 genes, also exhibited upregulation in FUS-sensitive neurons, but without distinction based on calcium response kinetics. This module was enriched in potassium channel-associated genes (e.g., Kcng2, Kcnip4, Kcnk13, Kcnmb1, and Kcnmb4) along with genes involved in cellular calcium ion homeostasis and pain response. The third module, containing 238 genes, was preferentially expressed in FUS-insensitive neurons and displayed enrichment in genes associated with collagen fibril organization and cell-matrix adhesion. This part of our study provides insights into the molecular determinants of FUS sensitivity and the diverse calcium response kinetics observed in the FUS responding neurons. It also highlights the involvement of specific gene modules associated with cellular processes crucial for neuronal function and response to external stimuli like FUS. Our sc-RNA-seq data also showed that among the FUS-sensitive neurons, more than half belonged to GINIP^+^ neurons. This was confirmed using GINIP^mCherry^ mice (10), in which we showed that 72% of mCherry^+^ neurons and only 47% of mCherry^-^ neurons responded to FUS stimulation. Using mCherry^+^ neurons in combination with IB4 binding, we found that 84% of C-LTMRs and 67% of MRGPRD-expressing C-HTMRs responded to FUS stimulation. Both neuronal subtypes have been shown to play critical roles in mechanosensation, with MRGPRD-expressing neurons being involved in mechano-nociception and C-LTMRs having dual functions: the sensation of pleasant touch under physiological conditions and modulation of mechanical pain under pathological conditions (21). Our study highlights the potential of using FUS as a non-invasive method to selectively target and activate specific neuronal subtypes within the DRG. The preferential activation of GINIP^+^ neurons, particularly C-LTMRs and MRGPRD-expressing C-HTMRs, suggests a nuanced approach to sensory pathway modulation that could be exploited for therapeutic benefit. Detailed investigations into the FUS parameters that can selectively activate different neuronal subtypes will be crucial to open up exciting possibilities for the development of innovative pain management therapies, highlighting the importance of targeted neuronal modulation using non-invasive techniques such as FUS.

## MATERIALS AND METHODS

### Mice

Mice were maintained under standard housing conditions (22°C, 40% humidity, 12h-light cycles, and free access to food and water). Special efforts were made to minimize the number of mice used in this study. All experiments were conducted on 6- to 12-weeks-old adult mice in accordance with European guidelines for the care and use of laboratory animals (Council Directive 86/609/EEC). Male AdvillinCre^Cre/+^ mice and female Ginip^fl/fl^ mice were bred. Only the Ginip^fl/+^::AdvillinCre^Cre/+^ mice were used for the study. Male AdvillinCre^Cre/+^ mice and female R26^GCamp6S/GCamp6S^ mice were bred. Only the R26^GCamp6S/+^::AdvillinCre^Cre/+^ mice were used for the study. AdvillinCre mice were generously provided by Dr Fan Wang (Duke University).

### Primary neuron preparation

All dorsal root ganglion (DRG) neurons were carefully extracted and digested with 2mg/mL of collagenase type II, 5mg/mL of dispase and 5mM of CaCl_2_ in HBSS media (magnesium- and calcium-free) containing 5mM HEPES, 10mM D-Glucose and 1% penicillin/streptomycin, at pH 7.5 for 1h at 37°C. After removing the supernatant, DRGs were resuspended with Leibovitz-15 complete medium (L-15 supplemented with 5% FCS and 1% penicillin/streptomycin), and then mechanically dissociated with two glass Pasteur pipettes of decreasing diameter (1 and 0.5 mm). The resulting suspensions were subjected to a density gradient centrifugation through Percoll (onto 12.5 and 28% Percoll gradient in Leibovitz-15 complete medium) at a speed of 1300g (acc=6, break=3) for 20 minutes, to eliminate cell debris. Cells were washed with Leibovitz-15 complete medium and centrifuged at 900g (acc=7, break=8) for 5 minutes. Cells were then resuspended in Neurobasal complete medium (Neurobasal A medium supplemented with 2mM L-glutamine, 1% penicillin/streptomycin and B27 1X) containing the following factors: Nerve growth factor (NGF) at 50ng/mL, Glial cell-derived neurotrophic factor (GDNF) at 50ng/mL, Neurotrophin-3 (NT3) at 50ng/mL, Neurotrophin-4 (NT4) at 10ng/mL and Brain Derived Neurotrophic Factor (BDNF) at 5ng/mL. Cells were plated in a Poly-D-lysine- (50 µg/ml) Laminin- (10 µg/ml) coated ibidi Polymer Coverslip Bottom (ibidi μ−Dish 35 mm, high-81156) and incubated at 37°C for 24 hours.

### Hydrophone measurements

The beam patterns in the elevation direction (X-Y) and azimuth plane (Y-Z) of the transducer were measured with a pressure measurement system composed of a 40 *µ*m-diameter needle hydrophone (Model NH0040, Precision Acoustics, UK), a preamplifier and a DC coupler. The hydrophone was fixed onto a three-axis motorized translation stage (M-403.4PD, Physik Instrumente, Germany), used to control the hydrophone position in the X-Y and Y-Z planes with a spatial resolution of 0.02 *µ*m. For each spatial position, the output signals of the DC coupler were acquired using a digitizer (PCI digitizer CS11G8, GaGe, Canada), in order to acquire the hydrophone signals at a sampling frequency of 250 MHz. The pressure maps were recorded in the X-Y plane (0.6 mm by 0.6 mm) with a step size of 0.01 mm, and in the Y-Z plane (3.8 mm by 0.8 mm) with a step size of 0.01 mm in the Y-direction and 0.02 mm in the Z-direction (Fig. 1*A* and *SI Appendix*, Figure S5*A*). The lateral and axial beam widths were measured to be 0.22 mm and 2.78 mm respectively at -6 dB.

### Calcium imaging and ultrasound stimulation setup

Calcium imaging experiments were performed on R26^GCamp6S/+^::AdvillinCre^Cre/+^ and Ginip^fl/+^::AdvillinCre^Cre/+^ mice. For Ginip^fl/+^::AdvillinCre^Cre/+^ mice, DRG neurons were incubated with 5 μM calcium dye (Fluo4-AM resuspended in DMSO) diluted in Opti-MEM solution for 45 minutes before the experiment. For every calcium imaging experiment, the culture medium was replaced by a Na-HEPES solution (5 mM KCl, 140 mM NaCl, 10 mM HEPES, 2 mM CaCl_2_, 2 mM MgCl_2_). Prior to FUS stimulation, the US beam was positioned at the center of the optical microscope’s field of view. To achieve this, the transducer was first operated in pulse-echo mode by using a pulser-receiver (5073PR, Olympus, France) and focused onto the culture dish containing an isolated polystyrene microbead with a radius of 20 *µ*m located at the center of the optical field of view. The transducer was moved laterally to examine and maximize the ultrasonic echo signal from the microbead, the echo signal being maximal when the microbead is at the center of the US beam focus. Calcium imaging experiments were performed using an inverted microscope (Olympus IX73, 20X objective with a numerical aperture of 0.45) equipped with an LED illuminator (pE-300white). Images were acquired at a sampling rate of 10 Hz and recorded with a Basler acA4096 camera. Primary neuronal cultures were performed as described above and stimulated with a FUS transducer (Olympus V317-SM, central frequency of 20 MHz, active diameter of 6.32 mm, focal distance of 12.7 mm). The FUS transducer was tilted at an angle of approximately 20° relative to the axis perpendicular to the culture dish in order to reduce standing wave formation within the cavity formed by the transducer and the culture dish (Fig. 1*A*). In addition, an acoustic absorber (Aptflex F28, Precision Acoustics, UK) was attached to the transducer to avoid reflected waves at the water/air interface. A function generator (33600B, Agilent, France) amplified with a power amplifier (75A250A, Amplifier Research, USA) drove the US transducer. The FUS stimulus consisted of a 20-MHz sinusoidal signal with peak positive pressure of 4 to 5 MPa and stimulus duration of 0.1 to 1 ms. Calcium responses were recorded before, during and after FUS stimulation. All experiments were performed at room temperature. Data were analyzed with an in-house MATLAB code (The Mathworks Inc., Natick, MA, USA). Data are presented as the relative fluorescence change normalized to the initial basal fluorescence (Δ*F*/*F_0_*). Around 140 to 220 cell bodies were analyzed per experiment. This corresponds to cultures of primary DRG neurons from two mice distributed in three Ibidi imaging dishes. For each Ibidi imaging dish, six different Region-Of-Interests (ROIs), corresponding to the -6 dB beam area, were stimulated by FUS. On average, 10 cell bodies are located within each ROI. For each US stimulus (specific time and acoustic pressure), the experiments were repeated on at least two different days.

The kinetics of the calcium responses were classified in three groups: fast, intermediate and slow. To this end, for each DRG neuron, the normalized amplitude calcium response was computed and the peak response duration, defined as the duration over which the amplitude of the response remained higher than 90% of the maximum amplitude, was determined. Fast, intermediate and slow kinetics were arbitrarily defined as peak response durations of less than 18 s, comprised between 18 and 70 s and of more than 70 s, respectively. The slow-kinetics responses correspond to calcium responses that never return to their initial basal fluorescence.

### Single-cell library preparation

After calcium imaging recording, the cytoplasmic content of the neuron was harvested using a glass pipette, transferred into a lysis solution (RNaseOUT 8 U.µl^-1^, 2% Triton X-100) and flash frozen. Reverse transcription was performed using a SMART-seq v4 Low Input Kit (Clontech) directly on the cell lysate according to the manufacturer’s protocol. After cDNAds amplification, 50 μl of the sample were subjected to cDNA purification on AMPure XP beads. 0.5 ng of each purified cDNAds were used to construct the sequencing library using the Nextera XT DNA Library Preparation Kit (Illumina) according the manufacturers protocol using fragments over 300-bp length from each neuron to generate single-cell cDNA libraries for mRNA sequencing.

### Processing, quality control and filtering of single-cell RNA-seq data

Sequencing was performed by the GenomEast platform, a member of the “France Genomique” consortium (ANR-10-INBS-0009). Single-cell libraries were sequenced on an Illumina HiSeq 4000 sequencer (Illumina) as single read 50 base reads with an average depth of 4.7 million reads per cell. Image analysis and base calling were performed using RTA version 2.7.7. Demultiplexing was performed using bcl2fastq version 2.20.0.422 and quality control was performed using FastQC v0.11.5n (Andrew, 2010). Bowtie v1.2.2 was used by FastQScreen v0.11.3 to map the sequences to the mouse genome (mm10 / GRCm38.90). Reads mapping to gene exons were counted using featureCounts57 (C version 1.4.6-p2). Read counts were analyzed in the R/Bioconductor environment (version 4.2.2; www.bioconductor.org). Genes with less than one read per cell on average were removed (n = 26,546 genes remaining) and the count data were normalized between samples by TMM (trimmed mean of the M-values) using the scran package in R Bioconductor (22). Quality control plots were performed using scran as described in Lun et al. (22). Relative expression levels were obtained by sample normalization using TPM (transcripts per kilobase million). Downstream analysis was performed using custom R scripts.

### Dorsal root ganglion neuron clustering

The mapping of the scRNAseq neurons in DRG neuron transcriptional clusters was done by a cross-dataset normalization approach. We used fastMNN, the new implementation of Mutual Nearest Neighbours (MNN) correction (8) implemented in the R package batchelor to combine our dataset with Zeisel and collaborators scRNAseq sensory neuron dataset (9). Briefly, we selected 3000 most variables genes from the Zeisel et al. data set, and found that 1091 of these genes were present among the 26,546 that we selected in our own data set. We then used this set of 1091 genes for the mapping of our cells to the reference data. A principal component analysis (PCA) was performed between Zeisel dataset (9) and our dataset to correct for batch effect. Identification of MNN was done in this reduced dimension space using the following parameters: dimension = 50 and k = 2 and 150 for our and Zeisel datasets, respectively. To assign each individual neuron to a specific cluster, we built a k nearest-neighbors (kNN) graph with k = 50. Individual neurons were assigned to one of the 17 clusters described in Zeisel et al. (PSNF1-3, PSNP1-6, PSPEP1-8). The nomenclature of Usoskin et al. (23) was preferred for clarity in the main figure (see *SI Appendix*, Fig. S2 for correspondence). Corrected expression values obtained during this procedure were only used for cluster assignment and visualization with t-distributed stochastic neighbor embedding (tSNE) projection (perplexity = 50, max iteration = 1000, theta = 0.35).

A second approach was used to assign each of our picked cells to one of the reference clusters. We log-transformed all counts from Zeisel et al. (9) with log2(x+1) transformation and averaged the log-transformed counts across all cells in each of the 17 clusters to obtain reference transcriptomic profiles of each cluster, using the same 1091 genes as above (17 × 1091 matrix). We applied the same log2(x+1) transformation to the TMM counts of our picked cells, and for each cell, computed Pearson correlations across the 1091 genes with all 17 Zeisel et al. (2018) clusters. Each cell was assigned to the cluster to which it had the highest correlation (Unweighted Paired Group Mean Arithmetic classifier) (*SI Appendix*, Fig. S2).

### Differential gene expression analysis

For differential gene expression analysis (DGE) data were analyzed using DESeq2 (24) and edgeR packages, and results were filtered for log2FC (fold change) and false discovery rate (FDR): |log2FC| > 1 and FDR < 10%. Given the known fact that analysis results differ among algorithms used (25, 26), differential expression of transcripts was only considered to be significant if determined so by both analysis algorithms (i.e., DESeq2 and edgeR) in line with recommended analysis procedures (25, 27). For the generation of heat maps shown in Figures 2 and 3, z-transformed TMM count data was used.

### Weighted gene co-expression network analysis

To perform WGCNA, a matrix of signed Pearson correlation between all pairs of transcripts was computed; This correlation matrix was raised to power β = 6 to calculate a adjacency matrix. To minimize the noise and spurious associations, the adjacency matrix was transformed to topological overlap matrix (TOM). The matrix 1-TOM was used as the input of average linkage hierarchical cluster, and genes with similar expression pattern were clustered together. The expression profile of a given gene module was represented by its first principal component (as known as module eigengene, ME) which can explain the most variation of the module expression levels. Module membership (also known as module eigengene based connectivity, kME) of each gene was calculated by correlated the gene expression profile with ME. The module labeled grey as a color corresponds to the set of genes which have not been clustered in any module. Gene ontology enrichment analysis was performed using Bioconductor package “topGO” (PMID:16606683). Terms were accepted if they are hitted more than 1 gene and Fisher exact Test P-value < 0.05.

### Modeling the relationship between gene expression and calcium response to ultrasound

We adapted the method developed by Bomkamp et al. (28). After accounting for the most variable genes among our dataset (∼14,000 genes), only genes which were expressed at a level of 1 TPM or higher in at least 80% of ultrasound-responsive neurons were included. Out of all genes represented in the RNA-seq dataset, 2398 passed this thresholding step. For the remaining genes, we fitted a linear model relating the maximum peak amplitude of the calcium response to ultrasound (P) to the expression of the gene (G) and cell class (C) using the R package stats v4.2.2. This model linked the calcium peak parameter to the gene and cell class (P∼G+C; “class-conditional model”).

### Deformation with high-speed imaging

Deformation experiments were performed on Ginip^fl/+^::AdvillinCre^Cre/+^ mice. To identify GINIP^+^ neurons, one image of the native fluorescence was recorded before each US stimulation. High speed imaging was performed using an inverted microscope (Olympus IX73, 20x objective with a numerical aperture of 0.45). Images were acquired at 100 000 fps and recorded with a Photron FASTCAM SA1.1 camera. The light source was a halogen cold light source (KL 1500 HAL, SCHOTT, France) manually positioned above the Ibidi imaging dish. The transducer was positioned to have the US beam at the center of the optical microscope’s field of view, as described above. Each neuron was stimulated several times consecutively: 6 consecutive times for the pressure series (5 MPa, 4 MPa, 4.2 MPa, 4.4 MPa, 4.6 MPa and 5 MPa) for a stimulus duration of 1 ms, or 7 consecutive times for the time series (1 ms, 0.1 ms, 0.2 ms, 0.4 ms, 0.6 ms, 0.8 ms, and 1 ms) for a peak positive pressure of 5 MPa. To ensure that consecutive FUS stimuli do not affect cell deformation, the optimal FUS stimulus (5 MPa, 1 ms) is repeated twice at the beginning and end of the series of stimuli, and neurons that did not exhibit the same projected area for these two optimal FUS stimuli were excluded from the analysis. Projected areas were recorded before, during and after US stimulation. Cells were imaged using Photron FASTCAM Viewer 4 software. All experiments were performed at room temperature. The deformability (D) was calculated as

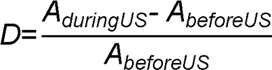

where *A_beforeUS_* and *A_duringUS_* represent the measured projected area of the cell before and during the FUS stimulation. To determine *A_duringUS_*, the maximum projected area was used, corresponding to the area on the last image of the stimulus duration. The area of the cells was manually quantified using an in-house MATLAB code (The Mathworks Inc., Natick, MA, USA), as shown in Fig. 4*C*.

Linear mixed models of increasing complexity were fitted to the deformation vs. acoustic pressure data using the R package stats (lme function), starting with a random-intercept model, and progressively adding the factors stimulus duration, neuron type (GINIP^+^ and GINIP^-^), and random slope. Models were compared using likelihood ratio tests (anova function). The same analysis procedure was repeated for the deformation vs. stimulus duration.

### Temperature elevation

The temperature rise induced by a US stimulus relates to the acoustic absorption coefficient of the medium insonified by the US wave. To compute this elevation theoretically, two media were considered: the polymer coverslip for which the acoustic absorption coefficient was measured to be 0.3 Np/cm/MHz, and DRG neurons for which the absorption coefficient was assumed to be close to the absorption value of the brain of 0.024 Np/cm/MHz^1.18^ (29). Following Fry and Fry (30), the temperature rise Δ*T* for a short exposure time is given by ΔT=QΔt/(Cρ_0_), where *Q* is the rate of heat generation per unit volume, Δ*t* is the time duration of the US stimulus, *C* and p_0_ are the specific heat capacity and density of the medium. The heat generation *Q* produced by US for a plane travelling wave is given by 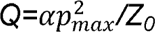, where *p_max_* is the maximum acoustic pressure inferred from hydrophone measurements, a is the absorption coefficient of the medium per unit path length and *Z_0_* is the acoustic impedance of the medium (31). The maximum temperature rise Δ*T*_max_ should be mainly due to the acoustic absorption coefficient of the polymer substrate, which is much higher than the absorption of the DRG neurons. We predicted that the US stimulus with maximum acoustic pressure of *p^+^*=3.6 MPa at 20 MHz for a pulse duration of Δ*t*=1 ms should produce a temperature rise on the polymer coverslip of 1.2°C at 20 MHz (and 0.5°C at 40 MHz when considering the maximum acoustic pressure of *p^+^*=1.6 MPa at this harmonic frequency; see *SI Appendix*, Table S1). We can therefore expect a temperature rise of less than 2°C. The following properties of the polymer coverslip, mainly composed of polyethylene, were considered: p_0_=1010 kg/m^3^, *Z_0_*=2.6 MRayl, *C*=2400 J/kg/K and a=6 Np/cm at 20 MHz (and 12 Np/cm at 40 MHz). In contrast, the temperature rise caused by the absorption of the DRG neurons is expected to be much smaller Δ*T_max-DRG_*= 0.15°C at 20°C by using the following parameters: *ρ_0_* =1020 kg/m^3^, *Z_0_* =1.54 MRayl, specific heat capacity approximated by that of water, *C*=4184 J/kg/K and a=0.82 Np/cm at the frequency of 20 MHz (29).

### Nonlinear simulations of the acoustic fields

Nonlinear simulations of the acoustic field produced by the 20-MHz transducer were conducted to confirm that the nonlinearity measured with the hydrophone was caused by the nonlinear propagation of the focused US.

The MATLAB’s k-Wave toolbox was used to simulate the acoustic field generated by our focused 20-MHz US transducer with focal distance of 12.7 mm and active element diameter of 6.35 mm. The simulations were performed in three dimensions on a domain of 8 x 8 x 18 mm^3^ with spatial discretization of 10 *µ*m. The Courant-Friedrichs-Lewy stability criterion was set to 0.15. The simulated stimulus waveform was a sinusoidal signal of 20 cycles at 20 MHz convolved with a gaussian window. The propagation medium was assumed to be water with the following characteristic properties: sound speed, *c* =1490 m/s; density, IZ =1000 kg/m^3^; absorption coefficient 2.17x10^-3^ dB/cm/MHz^2^; nonlinearity coefficient*, B/A*=5. The input pressure of the simulated transducer was empirically determined to obtain a peak positive pressure *p^+^* of either 0.5 MPa, 2.5 MPa or 5 MPa at the focus. Measured and simulated beam patterns in the Y-Z plane showed similar beam patterns at the fundamental (*SI Appendix*, Fig. S5*A*) and second harmonic (*SI Appendix*, Fig. S5*B*). The -6 dB transmit focal spot lengths determined by hydrophone measurements and k-Wave simulations were also in good agreement, both at the fundamental frequency (2.78 mm and 2.82 mm, respectively; *SI Appendix*, Fig. S5*C*) and at the second harmonic frequency (1.80 mm and 1.81 mm, respectively; *SI Appendix*, Fig. S5*D*). Nonlinear effects were evaluated in the spectral domain by estimating the amplitude of the second harmonic of the sinusoidal waves (*SI Appendix*, Fig. S5 *E* and *F*). The differences on the second harmonic amplitudes between experiments and simulations are less than 3 dB, which is satisfactory given the measurement uncertainties provided by the needle hydrophone manufacturer (16% and 24% of errors at 20 MHz and 40 MHz, respectively). Therefore, the nonlinear propagation of the focused US simulated here is sufficient to explain the nonlinear effects observed in experiments. Cavitation effects can be reasonably ruled out.

## Supporting information

Supplemental fiures

## Acknowledgments

We are grateful to the members of the two laboratories (the Moqrich lab at IBDM and the researchers at LMA) for the scientific discussions. The IBDM imaging and animal facilities for assistance. This work was funded by the French National Research Agency grant (ANR-HearUS) awarded to O.M, E.F and A.M and by the program Centuri to E.B. This work was also supported by institutional funding from the CNRS and Aix-Marseille Université to IBDM and LMA.

## Author Contributions

A.M, E.F and O.M designed the project, E.B and E.D performed the calcium imaging experiments and the FUS-mediated cell deformation, E.B performed the single cell RNA preparation and generated all the figures and managed the writing of the materials and methods section, T.P analysed the RNA-seq data and contributed to writing the corresponding chapter, S.Y performed the cell picking for the scRNA-seq, K.M and S.M performed the k-Wave simulations, P.M managed the mouse colonies used in this study, A.S performed the first RNA-seq analysis and provided input on the whole manuscript, A.M, E.F and O.M wrote the manuscript. All authors contributed to editing the manuscript.

