## Supplemental fiures for "Single cell RNA-sequencing reveals GINIP-expressing neurons as the main targets of focused ultrasound"

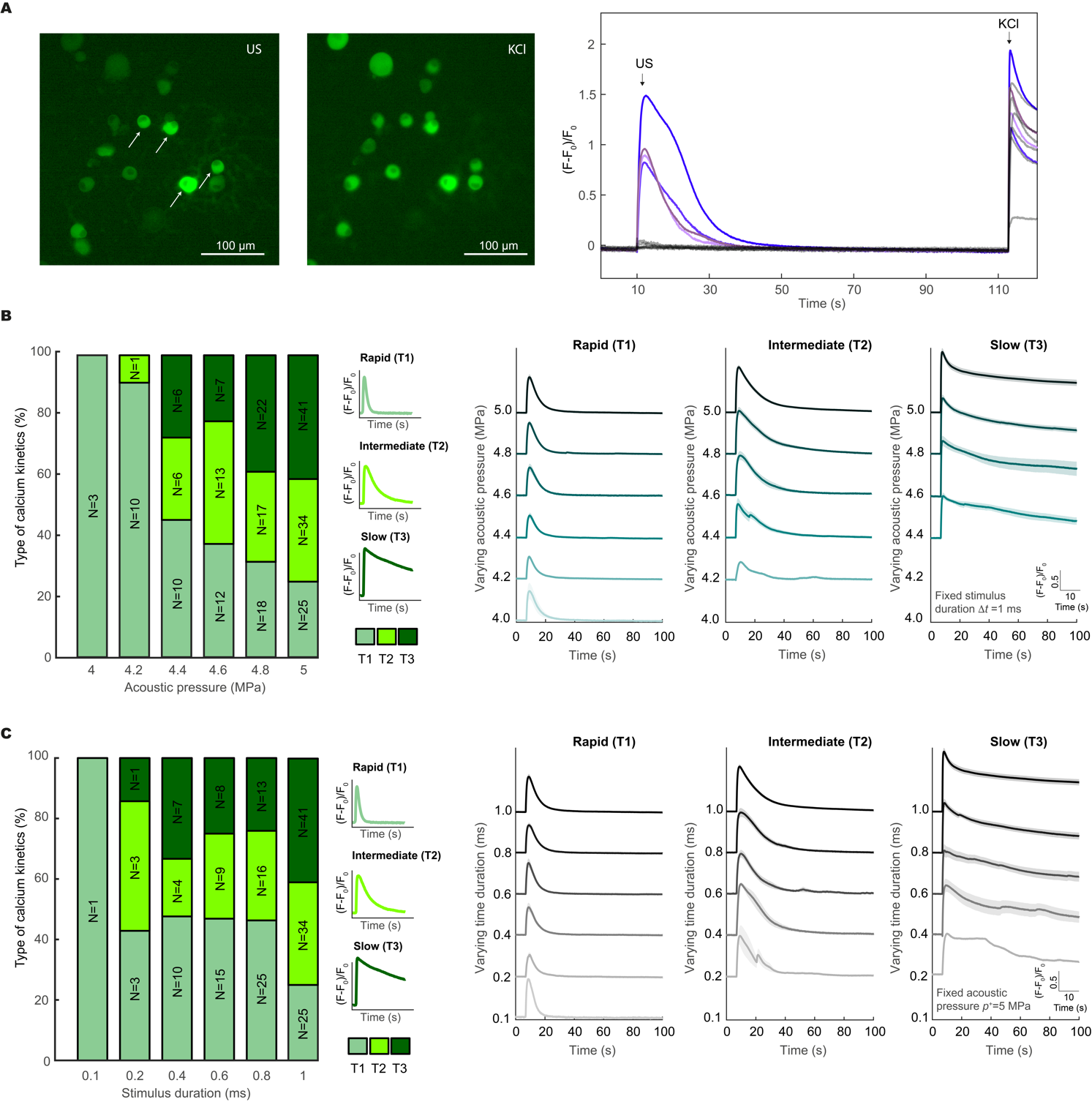

**Figure S1.** **Calcium kinetics of DRG neurons for varying acoustic pressures and stimulus durations.** (*A*) DRG neurons were stimulated by FUS at [*p^+^*=5 MPa, *Δt*=1 ms], followed by the application of potassium chloride (KCl, 100 mM) and analyzed by calcium imaging. All DRG neurons responded to KCl confirming the cell viability. (*B*) Quantification of the 3 types of calcium kinetics (slow, intermediate, rapid) observed in DRG neurons for varying acoustic pressures, and corresponding average calcium responses. N is the number of cells. The mean trace is shown in blue gradient and the SEM is shaded. (*C*) Quantification of the 3 types of calcium kinetics (slow, intermediate, rapid) observed in DRG neurons for varying stimulus durations, and corresponding average calcium responses. The mean trace is shown in grey gradient and the SEM is shaded. N is the number of cells.

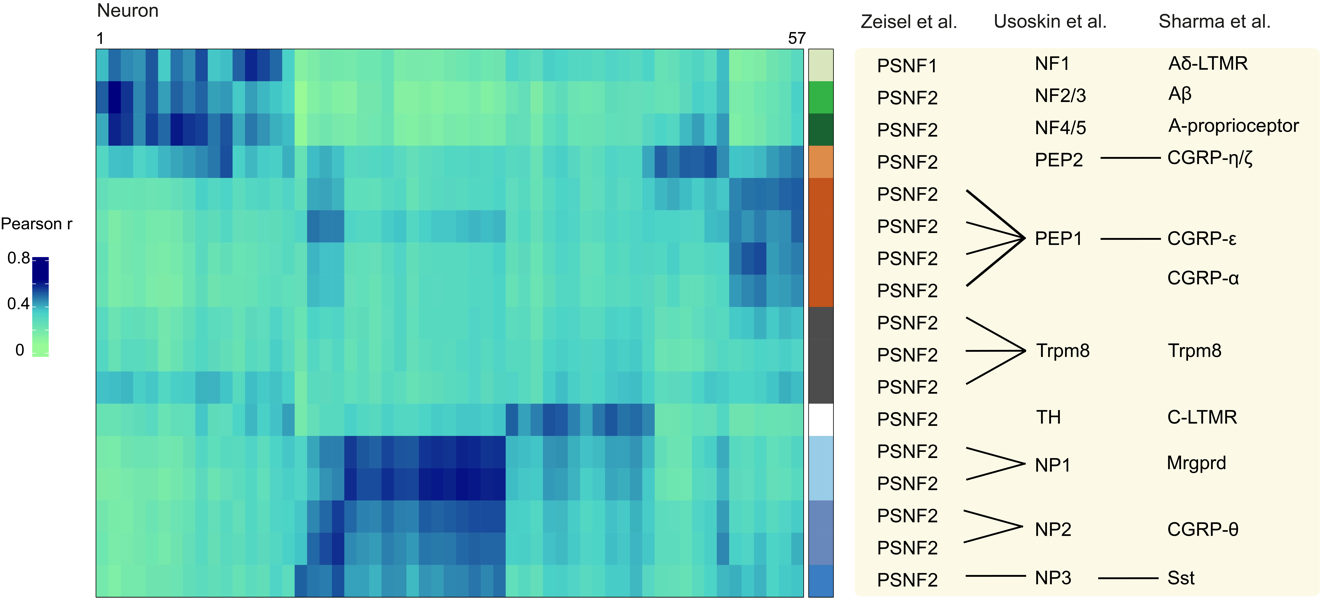

**Figure S2.** **Validation of the** **expression profiling of FUS-sensitive** **neurons.** Heatmap showing the correlation between each sequenced DRG neuron with each cluster of Zeisel et al. (9) using the averaged log-transformed counts of the 1098 genes used for the mapping.

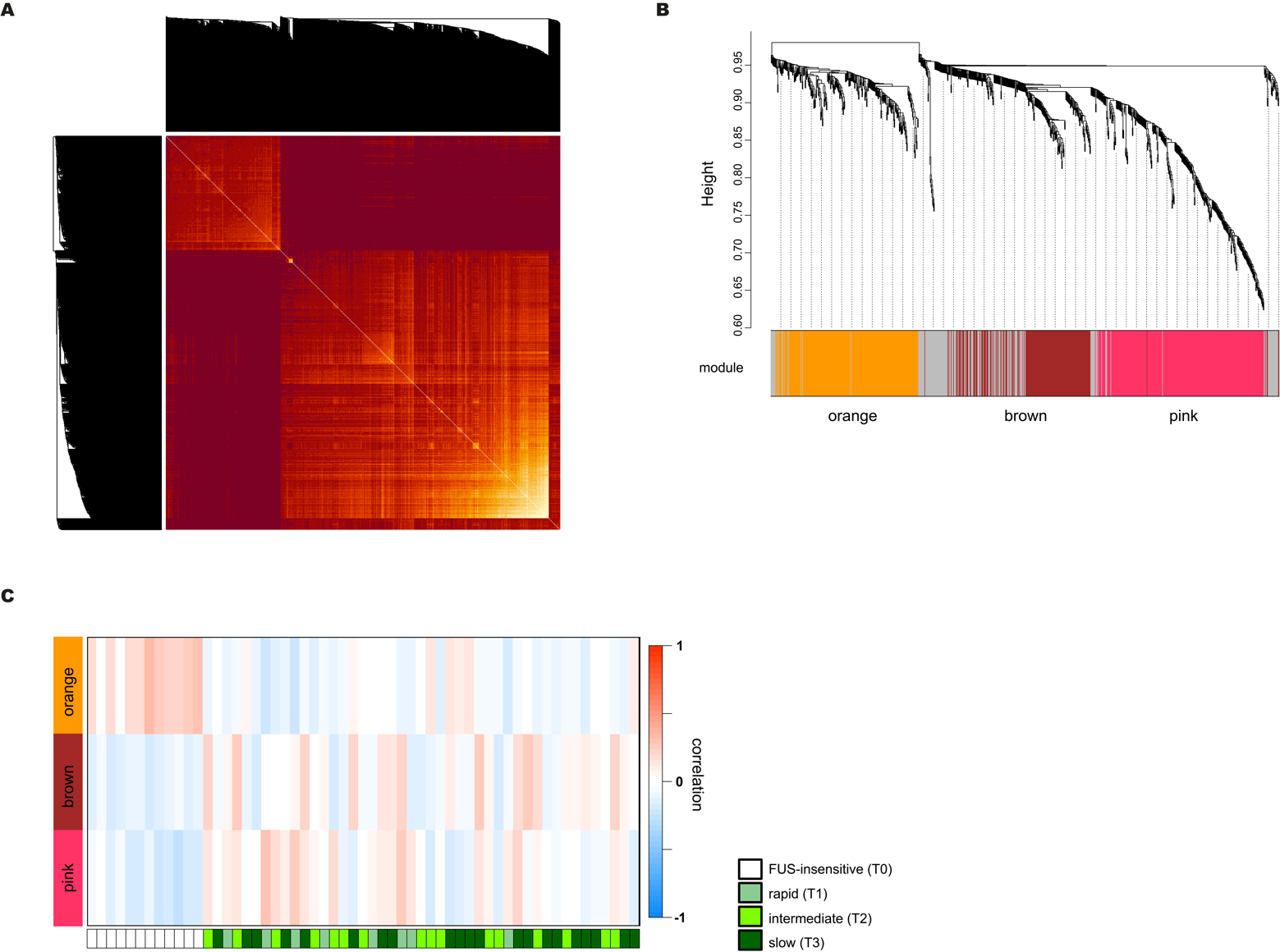

**Figure S3.** **Validation of the analysis of genes associated with the 3 types of calcium kinetics** (*A*) Weighted gene co-expression network analysis (WGCNA). Heatmap depicting the topological overlap matrix among all genes in the analysis. Blocks of lighter colors indicate the modules. (*B*) Hierarchical cluster tree showing co-expressed gene modules identified by WGCNA, modules correspond to each branch are labeled by colors. Grey module are unassigned genes. (*C*) Correlation matrix showing expression pattern of each 57 sequenced DRG neurons. Each cell contains Pearson correlations of gene expression profile from each neuron with each module eigengenes.

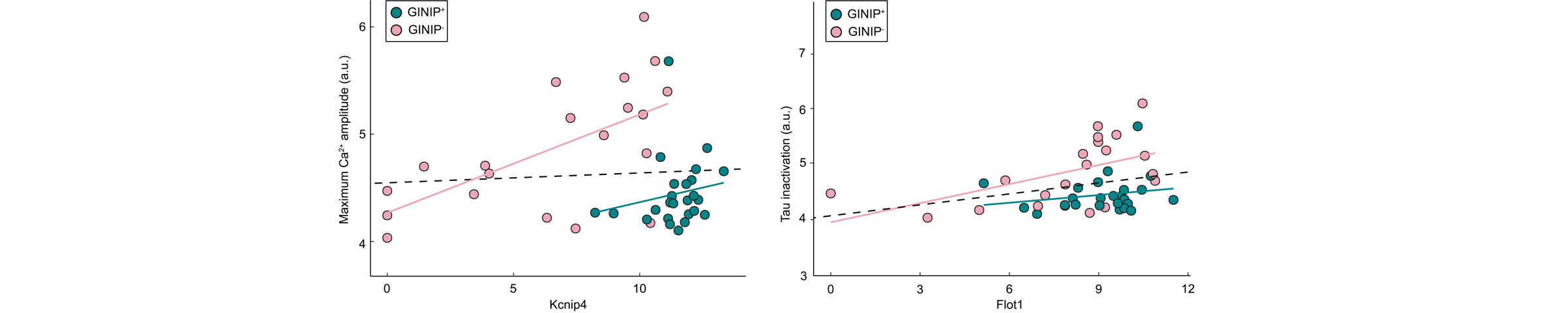

**Figure S4.** **Relationship between gene expression and amplitude of calcium response to FUS.** Example of genes showing significant association with the maximum calcium amplitude and inactivation kinetics that are class-conditional (*i.e.* significant after correction for cell class). Solid lines indicate linear fits within GINIP^+^ or GINIP^-^ neuron types, and dashed lines indicate linear fits including all cell types. Gene expression is quantified as TMM (Trimmed mean of M-value).

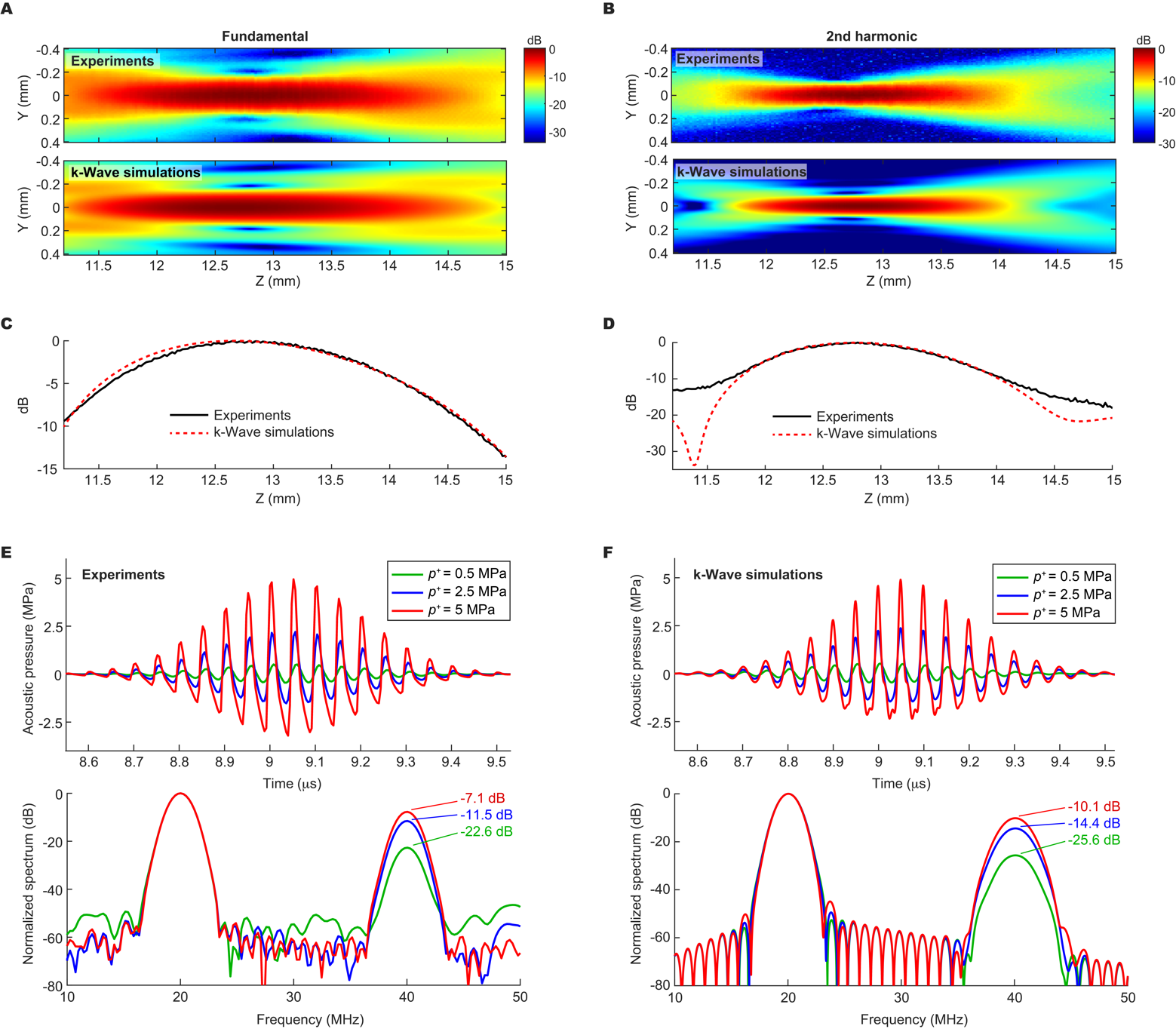

**Figure S5.** **Nonlinearity caused by the nonlinear propagation of the FUS.** (*A*) Hydrophone measurement and nonlinear k-Wave simulation of the FUS transducer’s pressure distribution in the Y-Z plane at the fundamental of 20 MHz and (*B*) at the second harmonic of 40 MHz.  (C) Measured and simulated axial beam profile at the fundamental of 20 MHz and (D) at the second harmonic of 40 MHz. (*E*) Sinusoidal temporal signals measured by the hydrophone at the focus and filtered by Gaussian and Tukey windows for three different peak positive pressures of 0.5, 2.5 and 5 MPa. The corresponding normalized spectra are also shown. (*F*) Simulated waveforms at the focus for three different peak positive pressures of 0.5, 2.5 and 5 MPa. The corresponding normalized spectra are also shown.

**Table S1.** Correspondence table between several ultrasound parameters deduced from the hydrophone measurements: the peak positive pressure *p^+^*, the peak negative pressure *p^-^* and the Spatial Peak Pulse Average intensity *I_sppa_* by considering all frequency content, as well as the peak positive pressure *p^+^* at the fundamental (20 MHz) and second harmonic (40 MHz).

| *p^+^*  (MPa) | *p^-^*  (MPa) | *I_sppa_*  (W/cm^2^) | *p^+^* at 20 MHz  (MPa) | *p^+^* at 40 MHz  (MPa) |
| --- | --- | --- | --- | --- |
| 4.0 | -2.6 | 346 | 3.0 | 1.1 |
| 4.2 | -2.7 | 380 | 3.1 | 1.2 |
| 4.4 | -2.8 | 416 | 3.2 | 1.3 |
| 4.6 | -3.0 | 453 | 3.4 | 1.4 |
| 4.8 | -3.1 | 494 | 3.5 | 1.5 |
| 5.0 | -3.2 | 536 | 3.6 | 1.6 |

**Supplementary dataset in the Excel file**

**Table E1.** Sample table of the DRG cells collected for scRNA sequencing after FUS stimulation. (see Excel file)

**Table E2.** Differentially expressed genes detected from DRG cell after FUS stimulation using R packages DEseq2 and edgeR together with fold change (log2TMM) and adjusted P-value.

(see Excel file)

**Table E3.** Gene list of each significant WGCNA modules.

(see Excel file)

**Table E4.** List of significant GO term associated with each WGCNA modules.

(see Excel file)

**Table E5.** List of genes significantly correlated with maximum amplitude of the calcium response or the constant of inactivation. The *p*-value<0,05 and *R²* > 0,20 were listed.

(see Excel file)
